# Epigenetic therapy to enhance therapeutic effects of PD-1 inhibition in uveal melanoma

**DOI:** 10.1101/2021.02.04.429575

**Authors:** Vasu R Sah, Henrik Jespersen, Joakim Karlsson, Mattias F Lindberg, Lisa M Nilsson, Lars Ny, Jonas A Nilsson

**Author notes:** Corresponding authors: Jonas A Nilsson, Sahlgrenska Center for Cancer Research, Box 425, University of Gothenburg, 40530 Gothenburg, Sweden. Harry Perkins Institute of Medical Research, 6 Verdun St, WA 6009, Nedlands, Perth, Australia.

## Abstract

Targeted therapy and immunotherapy have revolutionized the treatment of metastatic skin melanoma but none of the treatments are approved for patients with metastatic uveal melanoma (UM). Here we hypothesized that the poor responses to immunotherapy of UM can be enhanced by epigenetic modulation using HDAC or BET inhibitors (BETi). Cultured uveal melanoma cells were treated with the HDAC inhibitor (HDACi) entinostat or BETi JQ1. Entinostat induced HLA expression and PD-L1, but JQ1 did not. A syngenic mouse model carrying B16-F10 melanoma cells were treated with PD-1 and CTLA-4 inhibitors, which was curative. Co-treatment with the bioavailable BETi iBET-726 impaired the immunotherapy effect. Monotherapy of a B16-F10 mouse model with anti-PD-1 resulted in a moderate therapeutic effect that could be enhanced by entinostat. Mice carrying PD-L1 knockout B16-F10 cells were also sensitive to entinostat. This suggests HDAC inhibition and immunotherapy could work in concert. Indeed, co-cultures of UM with HLA-matched melanoma-specific tumor-infiltrating lymphocytes (TILs) resulted in higher TIL-mediated melanoma killing when entinostat was added. Further exploration of combined immunotherapy and epigenetic therapy in metastatic UM is warranted.

## Introduction

Uveal melanoma (UM) is a rare form of melanoma, with an incidence of approximately eight new cases per million per year in Sweden [1]. UMs originate from choroid, ciliary body, or iris melanocytes and are clinically and biologically different to cutaneous melanoma [2, 3]. The primary disease can in most cases be successfully treated with radiotherapy or enucleation, but almost one half of patients subsequently develop metastatic disease, usually to the liver [4, 5]. While targeted therapies and immune checkpoint inhibitors have revolutionized the treatment of metastatic cutaneous melanoma [6–8], there are still no effective treatments for patients with metastatic UM, who have a median survival of less than 12 months [9].

UM harbors oncogenic mutations in the genes encoding the G-protein-alpha proteins *GNAQ* or the mutually exclusive *GNA11, PLCB4* or *CYSLTR2,* and poor prognosis is associated with monosomy of chromosome 3 (Chr. 3) and inactivating mutations of the *BAP1* tumor suppressor gene [10–13]. Therefore, BRAF inhibitors frequently used in skin melanoma do not work in UM. Outcomes with immune checkpoint inhibitor monotherapy have been disappointing, with response rates typically below 5% [14, 15]. Despite this, there appears to be some level of immunity against UM, since expanded and adoptively transferred tumorinfiltrating lymphocytes (TILs) have therapeutic clinical effects [13, 16]. Tebentafusp, a bispecific protein immunotherapy targeting CD3 and melanoma-specific gp100, has also shown activity in early-phase clinical studies [17], and combined PD-1 and CTLA4 immune checkpoint inhibition appears to be more effective than monotherapy, albeit not as effective as in cutaneous melanoma [18].

With the notable exception of iris melanomas, which display a UV damage mutational signature [13], most UM display low tumor mutational burden (TMB) [19]. Other factors that could mediate poor responses to immunotherapy could be poor antigen processing and presentation or immune suppressive tumor microenvironments [20–22], especially in the liver [23]. Drugs targeting epigenetic regulators such as histone deacetylases (HDACs), BET bromodomain proteins, and methyltransferases are showing promise as cancer therapies by reversing oncogene transcription and modifying the tumor microenvironment [24]. HDAC inhibitors (HDACi) block the effects of myeloid-derived suppressor cells (MDSCs) and regulatory T cells (Tregs) [25, 26]; they enhance the expression of cancer antigens silenced during immunoediting [27]; and/or they trigger DNA damage and cell death to activate danger signals and recruit immune cells [28, 29]. Finally, HDACi can increase HLA class I expression, resulting in enhanced antigen presentation [30].

The checkpoint ligand PD-L1 is usually induced when T cells meet cancer cells but HDACi can directly induce PD-L1 to inactivate T cells [31]. This is contrary to BET inhibitors (BETi) in some tumor types where PD-L1 is suppressed [32]. Nuclear acetylated PD-L1 was recently shown to stimulate antigen presentation [33], providing a potential explanation for why PD-L1-high tumors are sensitive to PD-1 inhibition. Since PD-L1 is induced by HDACi this suggests that anti-PD-1 therapies and HDACi could synergize. Previous *in vivo* preclinical studies [26, 31, 34, 35, 36–38] and phase I/II trials have shown encouraging results when combining the HDACi [39–42]. However, it is unknown whether this combination is effective in metastatic UM.

Here we investigate if HDACi or BETi increase UM immunogenicity (e.g., by inducing HLA-1), induces PD-L1, and thereby synergizes with immunotherapy in animal models.

## Results

### Entinostat alters the transcriptome of immune-related genes in UM cells

To assess the effect of HDAC inhibition on HLA and PD-L1 expression, the human UM cell lines 92-1 (mutations in *GNAQ* and *EIF1AX,* derived from a primary eye tumor), MEL202 (mutant *GNAQ* and *SF3B1,* primary tumor), MP41 (mutant *GNA11,* monosomy Chr. 3, primary tumor) and UM22 (mutant *GNAQ* and *BAP1,* metastasis) were treated with the HDAC inhibitor entinostat and analyzed by flow cytometry. Entinostat induced HLA-ABC in 92-1, MEL202, and UM22 UM cells, but HLA-ABC was already highly expressed in MP41 cells and not further induced (**Fig. 1a**, gating strategy shown in **Supplementary Fig. 1**). PD-L1 was induced by entinostat in all cell lines (**Fig. 1b**). To gain further insight into immune-related expression changes, gene expression changes following entinostat treatment were analyzed by RNA sequencing. This analysis confirmed induction of HLA genes and/or *CD274* (PD-L1) with RNA-seq for UM22, MP41, and 92-1 (**Fig. 1c**, **Supplementary Table 1**). Entinostat also induced the immune proteasome gene *PSMB9* and T cell cytokine genes *IL15* and *CXCL12* but not the ABC transporters *TAP1* and *TAP2.* Expression of the immune checkpoint protein TIM3 ligand *HMGB1* was suppressed in all cell lines and the ligand *CEACAM1* in all except UM22 (**Fig. 1c**, **Supplementary Fig. 2a,b**). These effects were not seen with the BET bromodomain inhibitor (BETi) JQ1 (**Fig. 1c**).

**Fig. 1.**
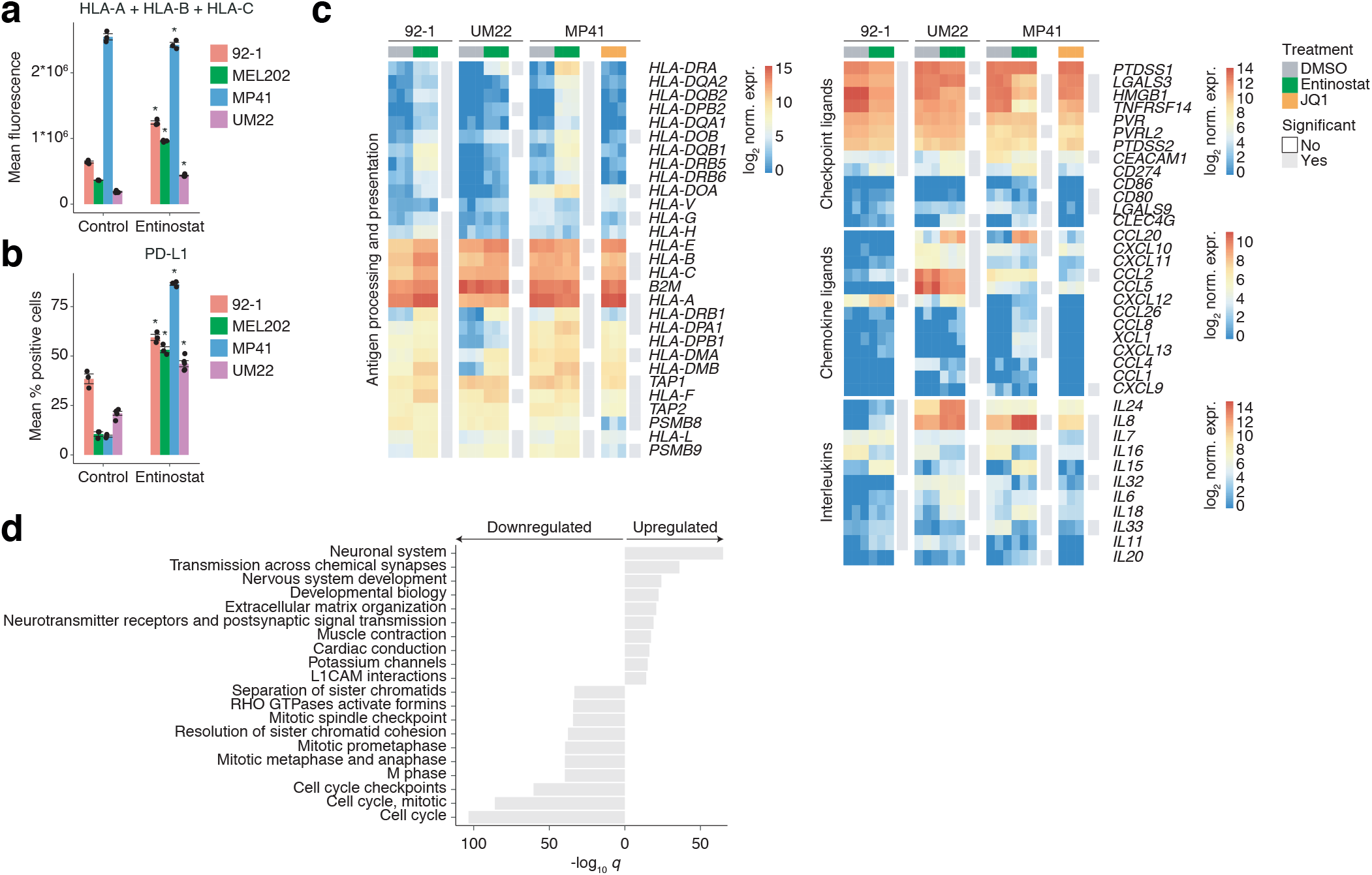
Entinostat regulates expression of immune-associated genes in human UM cell lines. **(a-b)** Human UM cell lines 92-1, MEL202, MP41, and UM22 were treated with DMSO or 1 μM entinostat for 48 h. Flow cytometry of (a) human HLA-ABC expression (mean fluorescence intensity **(b)** and human PD-L1 expression (% positive cells compared to unstained control). *n* = 3 biological replicates per cell line and condition were used, except for UM22, where *n* = 5 *and n* = 1 replicates were used. Significance was assessed with *t*-tests and adjusted *p*-values <0.05 were considered statistically significant, as indicated with asterisks. **(c)** Differentially expressed immune-associated genes in the human UM cell lines 92-1, MP41, and UM22 after treatment with entinostat for 48 h compared to DMSO (*n* =3 biological replicates per condition). Genes with FDR-adjusted *p*-values <0.05 were considered statistically significant. Statistical tests were carried out using DESeq2. Asterisks indicate genes significant in all three cell lines, whereas individual cell line-specific significance is indicated in gray next to each heatmap. **(d)** Enriched Reactome pathways among genes with adjusted *p*-values < 0.05 and absolute log2 fold change > 2 in all three cell lines, assessed with the MSigDB gene set enrichment analysis tool.

### Entinostat increases the anti-tumoral effects of T cells in vivo and in vitro

To assess the immune modulatory effect of HDACi and BETi in an immune competent and syngeneic mouse transplant model we used the B16-F10 murine melanoma cells. Although these tumors did not originate from the uvea of the eye, B16-F10 cells resemble UM in that they do not harbor classical cutaneous melanoma *BRAF, NRAS,* or *NF1* mutations and the TMB is low [43]. Entinostat induced surface expression of MHC class I and II and PD-L1 (**Fig. 2a**, **Supplementary Table 2**), similar to in human UM cells.

**Fig. 2.**
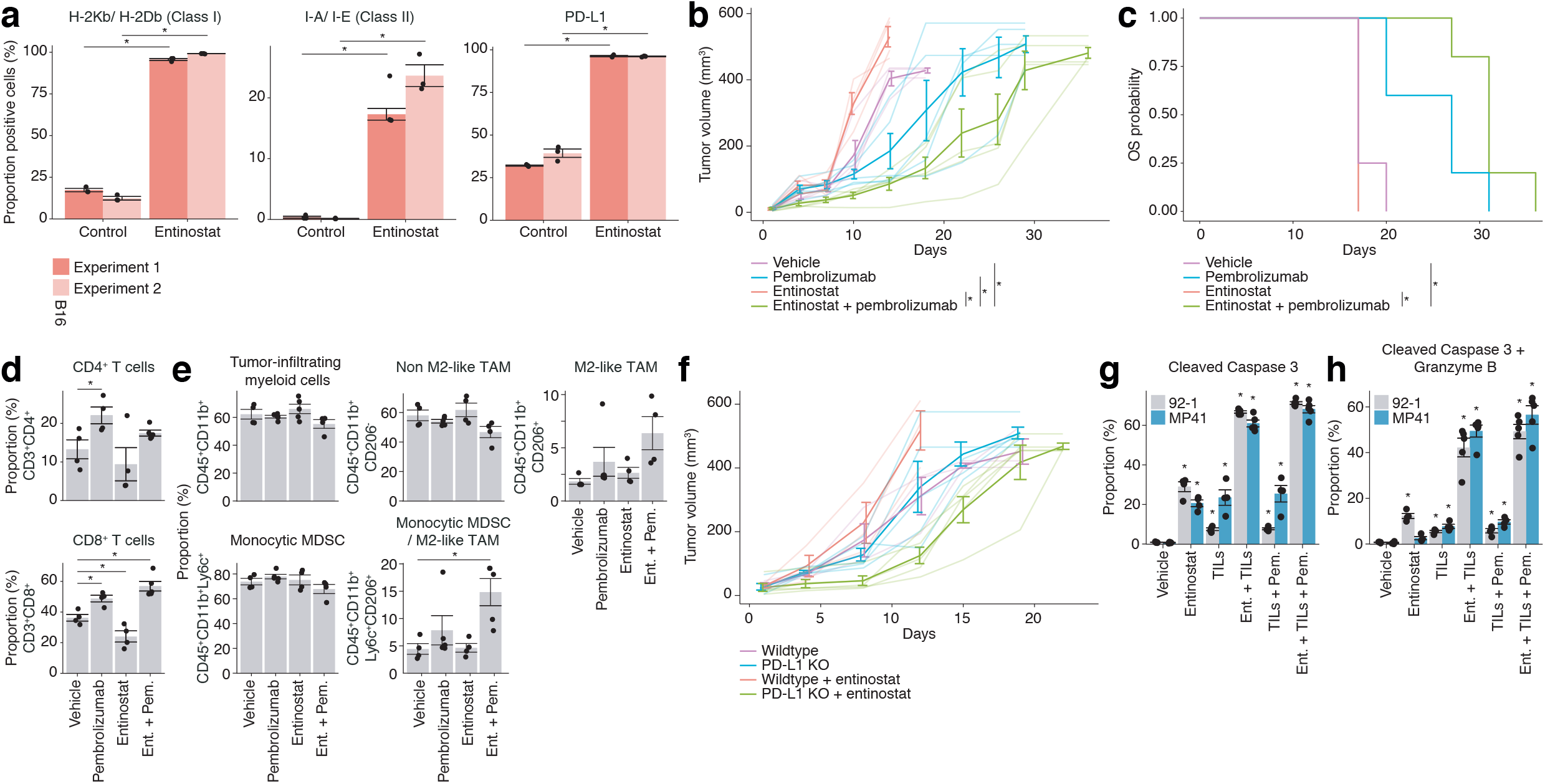
Entinostat enhances immunotherapy *in vitro* and *in vivo.* **(a)** Flow cytometry analysis showing HLA class 1, class 2 and PD-L1 expression in B16-F10 melanoma cells treated with entinostat. The experiment was repeated twice with *n* = 3 biological replicates each time. Asterisks indicate significance between vehicle and control. **(b-c)** Eighteen C57BL6 mice with subcutaneous B16-F10-luciferase tumors were allocated to groups to receive treatment with vehicle (*n*=4), entinostat (*n*=4), PD-1 inhibitor (*n*=5), or the combination of entinostat and PD-1 inhibitor (*n*=5). Tumors were measured with calipers and are plotted as mean volumes (bold lines) and individual volumes (light colored lines) (b). Asterisks indicate *p* < 0.05 as assessed with the “compareGrowthCurves” function in the *statmod* R package. Survival was plotted as a Kaplan-Meier curve (c). **(d-e)** End of study tumor samples from mice treated with indicated treatments were analyzed by flow cytometry to assess the distribution of tumor-infiltrating lymphocytes (d) and myeloid cells (e). For (d), *n* = 4 biological replicates were used per condition, except for the combination treatment, where *n* = 5 replicates were used. For (e), *n* = 4 biological replicates were used per condition, except for all treatments with pembrolizumab, treatments with entinostat in the experiment measuring CD45^+^CD11b^+^ cells, and treatment with entinostat + pembrolizumab in the experiment measuring CD45^+^CD11b^+^Ly6c^+^CD206^+^ cells, where *n* = 5 replicates were used. **(f)** Sixteen C57BL6 mice were injected subcutaneously with B16-F10-luciferase cells (*n*=6) or PD-L1-deficient CRISPR B16-F10-luciferase cells (*n*=10). Half of the animals in both groups received food containing entinostat. Tumors were measured with calipers and are plotted as mean volume (bold lines) and individual volumes (light colored lines). **(g-h)** HLA-A2:01-positive human UM cell lines 92-1 and MP41 were treated with DMSO, 1 μM entinostat, and 30 μg/ml pembrolizumab for 48 h with or without MART1-specific T cells for the last 24 h. Cells were fixed, permeabilized, and stained with antibodies targeting cleaved caspase-3 and granzyme B followed by flow cytometric analysis. Shown are the proportions of double-positive and single-positive melanoma cells. *n* = 4 biological replicates used per cell line and condition, except for assays with the combinations entinostat + TILs and entinostat + pembrolizumab + TILs, where *n* = 5 replicates were used. Significance of differences relative to vehicle (DMSO) were assessed with the two-tailed *t*-test and adjusted (Benjamini-Hochberg correction) *p*-values < 0.05 are indicated with an asterisk.

Next, we tested the *in vivo* efficacy of combined HDAC and PD-1 inhibition in C57/BL6 mice transplanted with subcutaneous B16-F10 tumors. Treatment with entinostat resulted in faster tumor growth than vehicle controls and PD-1 inhibitor alone did not inhibit tumor growth (**Fig. 2b,c**). However, combined entinostat and pembrolizumab significantly delayed tumor growth and prolonged survival compared to monotherapy (**Fig. 2b,c**). Combination treatment also increased intra-tumoral CD8^+^ T cells (but not CD4^+^ cells) and decreased both tumorinfiltrating myeloid cells and monocytic myeloid-derived suppressor cells (MDSCs). There was also a shift in macrophage phenotype, with increased proportions of pro-tumorigenic “M2-like” tumor-associated macrophages (TAMs) in combination therapy tumors (**Fig. 2d,e**).

CRISPR/Cas9 inactivation of *Cd274* (PD-L1) in implanted B16-F10 cells (**Supplementary Fig. S2c**) did not result in a slower tumor growth but it did ameliorate the faster growth induced by entinostat in parental B16-F10 cells. In fact, *Cd274* knockout cells grew slower than parental cells when treated with entinostat, consistent with the results from the pharmacological combination treatment (**Fig. 2f**). To investigate whether entinostat could impact on T cell killing of human UM cells, MART-1-specific T cells were isolated from an UM tumor using HLA-A2-specific MART-1 tetramers, expanded, and then used in killing assays. Incubation of HLA-A2-positive 92-1 and MP41 cells with MART-1-specific T cells induced UM cell apoptosis as measured by cleavage of caspase-3 and deposition of granzyme B (**Fig. 2g,h**). Addition of anti-PD-1 pembrolizumab moderately increased T cell killing.

Collectively, these data suggest that combined immune checkpoint blockade and HDAC inhibition can stimulate T cell immunity against human UM *in vitro* and *BRAF, NRAS,* and *NF1* wildtype melanoma *in vivo.*

### BET inhibition impairs immunotherapy in vivo

The finding that BETi JQ1 did not induce similar transcriptional changes as did entinostat (**Fig. 1c**) prompted further investigation into if BET inhibition would impact immunotherapy. Flow cytometry analysis of BETi-treated cells confirmed the RNAseq data and showed that HLA class 1 and 2 and PD-L1 expression was unchanged in UM22 cells and MP41 following treatment with JQ1 (**Fig. 3a**). In B16-F10 cells HLA class 1 was unchanged and PD-L1 was suppressed following JQ1 treatment (**Fig. 3b**), contrary to the effects of entinostat. To assess the negative impact of BET inhibition *in vivo* we treated B16-F10 melanoma bearing mice with anti-CTLA4 and anti-PD1 antibodies, to ensure better immunotherapy effects than by PD1 inhibition. Concomitant treatment with the bioavailable compound iBET726 resulted in a robust early response to treatment (**Fig. 3c,d**). Long-term the tumors grew back resulting in a worse survival of mice treated with combination BET inhibition and immunotherapy compared to immunotherapy alone (**Fig. 3e,f**). This suggests that although BET inhibition can work in monotherapy, it also inhibits immunotherapy with PD1/CTLA4 inhibitors.

**Fig. 3.**
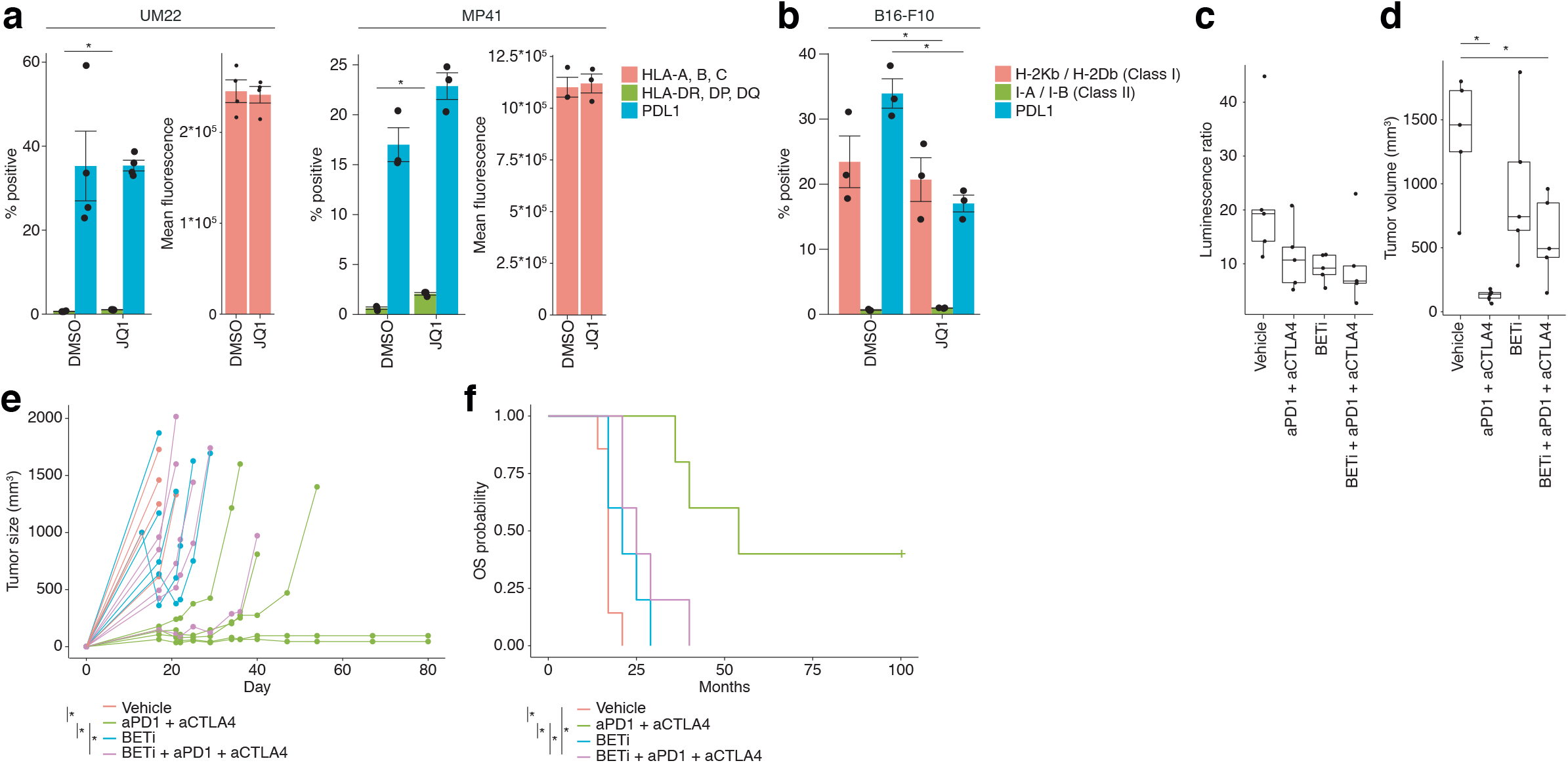
BET inhibition inhibits the expression of MHC class 1 and PD-L1 and the effect of immune checkpoint inhibition in vivo. (**a)** Flow cytometry of MHC class 1, MHC Class 2 and PD-L1 expression, in the uveal melanoma cell lines UM22 and MP41 treated with the vehicle DMSO or 1 μM of the BET inhibitor JQ1 for 48 hours. (**b**) Flow cytometry of MHC class 1 expression and PD-L1 expression in the mouse melanoma cell line B16-F10 treated with the vehicle DMSO or 1 μM of the BET inhibitor JQ1 for 48 hours. The experiments were repeated twice with *n* = 3 biological replicates for B16-F10, MP41 and for UM22, n=4 and n=2 replicates were analyzed. Asterisks indicate *p* < 0.05 with two-tailed t-tests. (**c-f**) Twenty C57BL6 mice with subcutaneous B16-F10-luciferase tumors were allocated in groups to receive treatment with vehicle (*n* = 5), CTLA4 + PD1 inhibitors (*n* = 5), iBET762 or combined iBET762 and CTLA4 + PD1 inhibitor. One week after treatment initiation, mice were imaged and luciferase activity was plotted (**c**). Tumors were also measured three weeks after treatment initiation (**d**) and followed until reaching ethics limit or up to 80 days post transplantation (**e**). In (e), asterisks indicate *p* < 0.05 with two-tailed t-tests. Survival was plotted as the time until the mice reached the ethics limit and were sacrificed (**f**). In (e) and (f) asterisks indicate adjusted *p*-values < 0.05, as assessed with the *compareGrowthCurves* function of the *statmod* R package in (e) and log-rank tests in (f).

## Discussion

Here we tested the hypothesis that epigenetic modulation can impact immunotherapy. Previous studies have shown that HDAC inhibitors modulate immune gene expression in cancer, including in HLA genes [30, 44]. However, as shown in other cancer types, and here in mouse melanoma *in vivo* and human UM *in vitro,* the trade-off is that entinostat monotherapy also induced PD-L1 in cancer cells. This may counteract any beneficial immunotherapeutic effects of HDAC inhibition. Indeed, entinostat-treated B16-F10 melanoma cells grew faster, an effect reversed on *Cd274* (PD-L1 gene) knockdown using CRISPR. This provided a strong rationale to combine HDAC and PD-1 inhibition to leverage the positive immune stimulatory effects of both drugs.

BETi have been deemed promising agents for treatment of cancer but a decade after the disclosure of JQ1, no drug has reached a phase III clinical trial. Their mechanism of action is clearly defined in vitro but problems with dose-limiting toxicities, efficacy and resistance have made progress slow thus far in patients. Some of these issues may also be due to the selection of indication as well, since BETi in parallel to development as anti-cancer drugs also show promise as anti-inflammatory drugs [45]. It may well be that the anti-tumoral effects of BETi are overridden by an inhibition of anti-tumoral immunity. Without powerful elimination of the BET-inhibited cancer cells by immune cells, treatment resistance may form. In the B16-F10 model used herein we observed that combined anti-PD1 and anti-CTLA4 treatment could result in durable responses in half of the treated mice but if they were also treated with BETi they quickly relapsed. This is in line with previous studies suggesting that BETi can inhibit priming by dendritic cells [46–48] as well as the proliferation [49] or function [50] of T cells. Also NK cell killing is suppressed by BETi via downregulation of NK cell ligands [51].

The above described data, and other published data showing that HDAC inhibition stimulates immunotherapy, have motivated us to initiate a clinical trial to test combined entinostat and pembrolizumab in patients with metastatic UM (NCT02697630, [52]). The data of this trial will be reported elsewhere.

## Methods

### Cell culture

B16-F10, a murine melanoma cell line, was obtained from Cell Lines Services (Eppelheim, Germany), while 92-1, MEL202 and MP-41, three human uveal cell lines, were obtained from the EACC and ATCC, respectively. UM22, a human UM cell line derived from a patient with UM [13], was grown in culture and used for further experiments. All cells were maintained in complete medium (RPMI-1640 supplemented with 10% FBS, glutamine, and gentamycin) and cultured at 37°C with 5% CO_2_. Cell line validation was performed by RNAseq where known and unique combinations of GNAQ/GNA11/SF3B1/EIF1AX/BAP1 driver mutations were confirmed.

To generate a Cd274 (PD-L1) CRISPR/Cas9 knockout B16-F10 cell line, Cas9:crRNA:tracrRNA ribonucleoprotein complex was assembled according to the manufacturer’s recommendations (Integrated DNA Technologies, Coralville, IA) and transfected into cells using Neon electroporation (Thermo Fisher Scientific, Waltham, MA). Negative cells were sorted for the absence of PD-L1 by staining with a PE-labeled anti-mouse PD-L1 antibody (clone MIH5, BD Biosciences, Franklin Lakes, NJ) using a FACSAria III (BD Biosciences). Absence of PD-L1 expression in the PD-L1 knockout cells was confirmed in cells treated with entinostat (Selleck Chemicals, Houston, TX) to induce PD-L1.

### Generation of MART-1 specific T cells

MART-1-specific T cells from uveal melanoma biopsies were identified as previously described (13) and sorted using FACSAria III (BD Biosciences). Sorted MART1-specific T cells were co-cultured with irradiated allogenic peripheral blood leukocytes at a 1:200 ratio in AIM-V cell culture medium (Invitrogen, Carlsbad, CA) supplemented with 6000 IU recombinant IL-2 (PeproTech, Rocky Hill, NJ), 10% human AB serum (Sigma Aldrich, St Louis, MO), and 30 ng/ml CD3 antibody (clone OKT3, Miltenyi Biotech, Bergisch Gladbach, Germany) for 14 days with regular media changes. After completion of the expansion protocol, MART1 specificity was confirmed using MART1-specific dextramers (Immudex, Copenhagen, Denmark).

### Animal experiments

All animal experiments were performed in accordance with EU Directives (regional animal ethics committee of Gothenburg #2021/19). Tumor models of parental B16-F10-luciferase or PD-L1-knockout B16-F10-luciferase cells were established by injecting 7.5 × 10^4^ cells per mouse mixed with an equal volume of Matrigel (Corning Inc., Corning, NY) subcutaneously into the flanks of four-to-six-week-old C57BL6 mice. Tumors were measured with calipers at regular intervals and tumor volumes calculated using the formula: tumor volume (mm^3^) = (length (mm)) × (width (mm) x width (mm))/2. Three days after transplantation, sedated mice were injected with 100 μl (30 mg/ml D-luciferin) in an isoflurane administrating chamber and then placed in an IVIS Lumina III XR machine (Perkin Elmer, Norwalk, CT). IVIS values on day three post tumor implantation were taken to allocate mice into balanced treatment groups of PBS-injected, 200 μg PD-1-blocking antibody-injected (clone RMP1-14, BioXCell, Lebanon, NH) intraperitoneally twice per week for three weeks, entinostat-treated (food containing 50 mg/kg entinostat), or a combination of PD-1-injected and entinostat-treated mice. For iBET immunotherapy combination, mice were treated with vehicle or iBET726 orally (10mg/kg) once daily for seven days, 250 μg PD-1 and CTLA-4 blocking (clone 9H10, BioXCell, Lebanon, NH) antibodies were injected intraperitoneally thrice per week for four weeks or a combination of PD-1 CTLA-4 antibodies with iBET762 were used.

### Cell staining and in vitro assays

Tumor cells were seeded and treated with entinostat (1 μM) or JQ1 (1 μM) for 48 hours and thereafter stained for 30 min at 4°C with specific antibodies for flow cytometry. The following anti-human antibodies were used for surface staining: FITC-labeled mouse anti-human HLA-DR, -DP, -DQ (Clone Tu39, BD Biosciences); PE-labeled mouse anti-human HLA-ABC (Clone G46-2.6, BD Biosciences); and APC-labeled mouse anti-human PD-L1 (clone 29E2A3, Biolegend, San Diego, CA). The following anti-mouse antibodies we used for surface staining: Alexa Fluor 647-labeled H-2Kb/H-2Db – MHC Class I (clone 28-8-6, Biolegend); PE-labeled I-A/I-E – MHC Class II (Clone M5/114.15.2, BD Biosciences), and PE-labeled PD-L1 (clone MIH5, BD Biosciences). Dead cells were excluded from the analysis by applying gating strategies.

Tumor cells were seeded in 24-well plates and treated with entinostat (1 μM), MART-1^+^ REP TILs in a 1:5 ratio with tumor cells, and 30 μg/ml pembrolizumab. 48 hours later, all cells were fixed and permeabilized using the Fixation/Permeabilization Solution Kit (554714, BD Biosciences) and then incubated with FITC-labeled rabbit anti-active caspase-3 (clone C92-605, BD Biosciences) and PE-labeled mouse anti-human granzyme B (clone GB11, BD Biosciences) antibodies for 30 minutes at 4°C. Flow cytometry data were acquired using BD Accuri C6 and BD Accuri C6 plus (BD Biosciences).

Tumor-bearing mice were sacrificed and single-cell suspensions were generated from tumors and spleens using mechanical dissociation before being passed through a 70 μm filter. Tumor suspensions were stained with 7-AAD live/dead stain (Miltenyi Biotec, Woking, UK), FITC-labeled CD3e (clone-145-2C11, BD Biosciences), PE-labeled CD4 (clone GK1.5, Biolegend), and APC-labeled CD8a (clone 53-6.7, BD Biosciences) for analysis of TILs. A seven-color myeloid panel with BUV395-labeled CD45 (clone 30-F11, BD Biosciences), Alexa Fluor 700-labeled F4/80 (clone BM8, BD Biosciences), brilliant violet 421-labeled Ly-6G (clone 1A8, Biolegend), PE/cyanine7-labeled Ly-6C (clone HK1.4, Biolegend), brilliant violet 605-labeled CD206 (MMR) (clone C068C2, Biolegend), BUV737-labeled CD11b (clone M1/70, BD Biosciences), and live/dead yellow stain (Thermo Fisher Scientific) was created for analysis of tumor samples. The proportions of tumor-infiltrating myeloid cells (CD45^+^CD11b^+^), monocytic MDSCs (CD45^+^CD11b^+^Ly6c^+^), “M2-like” TAMs (CD45^+^CD11b^+^CD206^+^), non “M2-like” TAMs (CD45^+^CD11b^+^CD206^-^), and Mo-MDSC ^+^M2-like TAMs^+^ (CD45^+^CD11b^+^Ly6c^+^CD206^+^) were acquired on a BD LSRII flow cytometer using FACSDiva software (BD Biosciences) for acquisition and compensation and then analyzed using FlowJo software.

### Statistical analysis

For flow cytometry measurements of HLA genes and PD-L1 in 92-1, MEL202, and MP41 cells, and independently for H-2Kb/H-2Db and I-A/I-E, unpaired two-tailed t-tests were carried out to assess effects of treatment with entinostat with the t.test function in R (v. 3.6.0, default parameters). Normality was assessed with Shapiro-Wilk tests, using the shapiro.test function in R. For differences in cell type proportions estimated by flow cytometry, as well as regarding proportions of cells with cleaved caspase 3 or granzyme B, unpaired two-sample t-tests were used. For analysis of tumor growth in *in vivo* experiments, the compareGrowthCurves function in the statmod R package (v. 1.4.32) with the parameter nsim=10^5^ was used. For survival analysis of *in vivo* experiments, log-rank tests were performed with the survdiff function from the survival R package (v. 3.2-7) with the parameter rho=0. *p*-values were adjusted for multiple testing with the Benjamini-Hochberg method. All statistical tests in this study were two-sided, and all error bars represent standard error of the mean, unless otherwise stated. A complete set of statistical tests in the study are present in **Supplementary Table 2**.

## Acknowledgments

Grant support came from Cancerfonden (to J.A.N), Familjen Erling Persson (to J.A.N), Knut and Alice Wallenberg Foundation (to J.A.N.), Vetenskapsrådet (to J.A.N.), Sjöbergstiftelsen (to J.A.N.), BioCARE Strategic grants (to J.A.N.), Lion’s Cancerfond Väst (to J.A.N.), Västra Götaland Regionen ALF grant (to J.A.N. and L.N), Assar Gabrielsson fond (to V.S.), Gustaf V Jubileumsklinikens forskningsfond to L.N and Wilhelm & Martina Lundgrens Vetenskapsfond (to J.K). We thank Carina Karlsson and Sofia Stenqvist for technical support.

## Supplementary Figure Legends

**Supplementary Fig. 1. Gating strategy for flow cytometry analyses. (a)** A gating strategy for excluding debris and choosing tumor cells based on high forward scatter (FSC) was employed and used for estimating levels of HLA-A, -B, and -C, PD-L1 and HLA-DP, -DQ and -DR in different cell lines. Experiments from entinostat-treated UM22 cells are shown as representative examples. **(b)** Granzyme B and cleaved caspase 3 measurements in cell lines cocultured with MART-1-reactive TILs. Experiments from MP41 are shown as representative examples. **(c)** Within the in vivo B16-F10 tumor suspension, leukocytes were identified by a low side scatter (SSC) and low forward scatter (FSC) with gates for estimating levels of live CD3+ cells for CD4+ and CD8+ TILs, as well as **(d)** gates for estimating levels of live CD45 cells for CD11b+, Ly6c+, Ly6g+, CD206+ myeloid infiltrating cells.

**Supplementary Fig. 2. Entinostat increases HLA expression in human UM cell lines and mouse B16-F10 melanomas. (a)** HLA class 2 expression as assessed by flow cytometry in human UM cell lines 92-1, MP41 and UM22 treated with DMSO or entinostat. The experiment was repeated twice with *n* = 3 biological replicates per cell line and condition, except in the case of UM22, where *n* = 5 and *n* = 1 replicates were used for the first and second experiments, respectively (excluded from statistical tests due to nearly absent expression in all cases). Significance was assessed by t-tests and adjusted p-values <0.05 (Benjamini-Hochberg correction) were considered statistically significant, as indicated by asterisks. **(b)** Immune-associated gene expression levels inferred from RNA sequencing data after entinostat treatment, relative to DMSO controls, as shown in Figure 1c. **(c)** Flow cytometry analysis of parental B16-F10 cells and CRISPR/Cas9-generated PD-L1 knockout B16-F10 cells after treatment with entinostat for 24 h.

## Notes

### Competing Interest Statement

The authors have declared no competing interest.

## References

1. Bergman L, Seregard S, Nilsson B et al. Incidence of uveal melanoma in Sweden from 1960 to 1998. Invest Ophthalmol Vis Sci 2002; 43: 2579–2583.

2. Damato B. Treatment of primary intraocular melanoma. Expert Rev Anticancer Ther 2006; 6: 493–506.

3. Jager MJ, Shields CL, Cebulla CM et al. Uveal melanoma. Nat Rev Dis Primers 2020; 6: 24.

4. Kujala E, Makitie T, Kivela T. Very long-term prognosis of patients with malignant uveal melanoma. Invest Ophthalmol Vis Sci 2003; 44: 4651–4659.

5. Diener-West M, Reynolds SM, Agugliaro DJ et al. Development of metastatic disease after enrollment in the COMS trials for treatment of choroidal melanoma: Collaborative Ocular Melanoma Study Group Report No. 26. Arch Ophthalmol 2005; 123: 1639–1643.

6. Robert C, Long GV, Brady B et al. Nivolumab in previously untreated melanoma without BRAF mutation. N Engl J Med 2015; 372: 320–330.

7. Robert C, Schachter J, Long GV et al. Pembrolizumab versus Ipilimumab in Advanced Melanoma. N Engl J Med 2015; 372: 2521–2532.

8. Larkin J, Chiarion-Sileni V, Gonzalez R et al. Five-Year Survival with Combined Nivolumab and Ipilimumab in Advanced Melanoma. N Engl J Med 2019; 381: 1535–1546.

9. Khoja L, Atenafu EG, Suciu S et al. Meta-Analysis in Metastatic Uveal Melanoma to Determine Progression-Free and Overall Survival Benchmarks: an International Rare Cancers Initiative (IRCI) Ocular Melanoma study. Ann Oncol 2019.

10. Van Raamsdonk CD, Bezrookove V, Green G et al. Frequent somatic mutations of GNAQ in uveal melanoma and blue naevi. Nature 2009; 457: 599–602.

11. Van Raamsdonk CD, Griewank KG, Crosby MB et al. Mutations in GNA11 in uveal melanoma. N Engl J Med 2010; 363: 2191–2199.

12. Robertson AG, Shih J, Yau C et al. Integrative Analysis Identifies Four Molecular and Clinical Subsets in Uveal Melanoma. Cancer Cell 2017; 32: 204–220 e215.

13. Karlsson J, Nilsson LM, Mitra S et al. Molecular profiling of driver events in metastatic uveal melanoma. Nat Commun 2020; 11: 1894.

14. Algazi AP, Tsai KK, Shoushtari AN et al. Clinical outcomes in metastatic uveal melanoma treated with PD-1 and PD-1L antibodies. Cancer 2016; 122: 3344–3353.

15. Mignard C, Deschamps Huvier A, Gillibert A et al. Efficacy of Immunotherapy in Patients with Metastatic Mucosal or Uveal Melanoma. J Oncol 2018; 2018: 1908065.

16. Chandran SS, Somerville RPT, Yang JC et al. Treatment of metastatic uveal melanoma with adoptive transfer of tumour-infiltrating lymphocytes: a single-centre, two-stage, single-arm, phase 2 study. Lancet Oncol 2017; 18: 792–802.

17. Middleton MR, McAlpine C, Woodcock VK et al. Tebentafusp, a TCR/anti-CD3 bispecific fusion protein targeting gp100, potently activated anti-tumor immune responses in patients with metastatic melanoma. Clin Cancer Res 2020.

18. Pelster MS, Gruschkus SK, Bassett R et al. Nivolumab and Ipilimumab in Metastatic Uveal Melanoma: Results From a Single-Arm Phase II Study. J Clin Oncol 2020; JCO2000605.

19. Royer-Bertrand B, Torsello M, Rimoldi D et al. Comprehensive Genetic Landscape of Uveal Melanoma by Whole-Genome Sequencing. The American Journal of Human Genetics 2016.

20. Whelchel JC, Farah SE, McLean IW, Burnier MN. Immunohistochemistry of infiltrating lymphocytes in uveal malignant melanoma. Invest Ophthalmol Vis Sci 1993; 34: 2603–2606.

21. Maat W, Ly LV, Jordanova ES et al. Monosomy of chromosome 3 and an inflammatory phenotype occur together in uveal melanoma. Invest Ophthalmol Vis Sci 2008; 49: 505–510.

22. Makitie T, Summanen P, Tarkkanen A, Kivela T. Tumor-infiltrating macrophages (CD68(+) cells) and prognosis in malignant uveal melanoma. Invest Ophthalmol Vis Sci 2001; 42: 1414–1421.

23. Yu J, Green MD, Li S et al. Liver metastasis restrains immunotherapy efficacy via macrophage-mediated T cell elimination. Nat Med 2021; 27: 152–164.

24. Topper MJ, Vaz M, Marrone KA et al. The emerging role of epigenetic therapeutics in immuno-oncology. Nat Rev Clin Oncol 2020; 17: 75–90.

25. Shen L, Pili R. Class I histone deacetylase inhibition is a novel mechanism to target regulatory T cells in immunotherapy. Oncoimmunology 2012; 1: 948–950.

26. Kim K, Skora AD, Li Z et al. Eradication of metastatic mouse cancers resistant to immune checkpoint blockade by suppression of myeloid-derived cells. Proc Natl Acad Sci U S A 2014; 111: 11774–11779.

27. Maio M, Coral S, Fratta E et al. Epigenetic targets for immune intervention in human malignancies. Oncogene 2003; 22: 6484–6488.

28. Landreville S, Agapova OA, Matatall KA et al. Histone Deacetylase Inhibitors Induce Growth Arrest and Differentiation in Uveal Melanoma. Clin Cancer Res 2012; 18: 408–416.

29. Lee JH, Choy ML, Ngo L et al. Histone deacetylase inhibitor induces DNA damage, which normal but not transformed cells can repair. Proceedings of the National Academy of Sciences of the United States of America 2010; 107: 14639–14644.

30. Campoli M, Ferrone S. HLA antigen changes in malignant cells: epigenetic mechanisms and biologic significance. Oncogene 2008; 27: 5869–5885.

31. Woods DM, Sodre AL, Villagra A et al. HDAC Inhibition Upregulates PD-1 Ligands in Melanoma and Augments Immunotherapy with PD-1 Blockade. Cancer Immunol Res 2015;3:1375–1385.

32. Zhu H, Bengsch F, Svoronos N et al. BET Bromodomain Inhibition Promotes Anti-tumor Immunity by Suppressing PD-L1 Expression. Cell Rep 2016; 16: 2829–2837.

33. Gao Y, Nihira NT, Bu X et al. Acetylation-dependent regulation of PD-L1 nuclear translocation dictates the efficacy of anti-PD-1 immunotherapy. Nat Cell Biol 2020.

34. Zheng H, Zhao W, Yan C et al. HDAC Inhibitors Enhance T-Cell Chemokine Expression and Augment Response to PD-1 Immunotherapy in Lung Adenocarcinoma. Clin Cancer Res 2016; 22: 4119–4132.

35. Kim YD, Park SM, Ha HC et al. HDAC Inhibitor, CG-745, Enhances the Anti-Cancer Effect of Anti-PD-1 Immune Checkpoint Inhibitor by Modulation of the Immune Microenvironment. J Cancer 2020; 11: 4059–4072.

36. Christmas BJ, Rafie CI, Hopkins AC et al. Entinostat Converts Immune-Resistant Breast and Pancreatic Cancers into Checkpoint-Responsive Tumors by Reprogramming Tumor-Infiltrating MDSCs. Cancer Immunol Res 2018; 6: 1561–1577.

37. Orillion A, Hashimoto A, Damayanti N et al. Entinostat Neutralizes Myeloid-Derived Suppressor Cells and Enhances the Antitumor Effect of PD-1 Inhibition in Murine Models of Lung and Renal Cell Carcinoma. Clin Cancer Res 2017; 23: 5187–5201.

38. Zimmer L, Livingstone E, Hassel JC et al. Adjuvant nivolumab plus ipilimumab or nivolumab monotherapy versus placebo in patients with resected stage IV melanoma with no evidence of disease (IMMUNED): a randomised, double-blind, placebo-controlled, phase 2 trial. Lancet 2020; 395: 1558–1568.

39. Sullivan RJ, Moschos SJ, Johnson ML et al. Abstract CT072: Efficacy and safety of entinostat (ENT) and pembrolizumab (PEMBRO) in patients with melanoma previously treated with anti-PD1 therapy. Cancer Res (13 Supplement) CT072; (79).

40. Gandhi L, Janne PA, Opyrchal M et al. Efficacy and safety of entinostat (ENT) and pembrolizumab (PEMBRO) in patients with non-small cell lung cancer (NSCLC) previously treated with anti-PD-(L)1 therapy. Journal of Clinical Oncology 2018; 36: 9036–9036.

41. Gray JE, Saltos A, Tanvetyanon T et al. Phase I/Ib Study of Pembrolizumab Plus Vorinostat in Advanced/Metastatic Non-Small Cell Lung Cancer. Clin Cancer Res 2019; 25: 6623–6632.

42. Rodriguez CP, Wu QV, Voutsinas J et al. A Phase II Trial of Pembrolizumab and Vorinostat in Recurrent Metastatic Head and Neck Squamous Cell Carcinomas and Salivary Gland Cancer. Clin Cancer Res 2020; 26: 837–845.

43. Castle JC, Kreiter S, Diekmann J et al. Exploiting the Mutanome for Tumor Vaccination. Cancer Res 2012.

44. Bhadury J, Nilsson LM, Muralidharan SV et al. BET and HDAC inhibitors induce similar genes and biological effects and synergize to kill in Myc-induced murine lymphoma. Proc Natl Acad Sci U S A 2014; 111: E2721–2730.

45. Huang D, Rossini E, Steiner S, Caflisch A. Structured water molecules in the binding site of bromodomains can be displaced by cosolvent. ChemMedChem 2014; 9: 573–579.

46. Remke N, Bisht S, Oberbeck S et al. Selective BET-bromodomain inhibition by JQ1 suppresses dendritic cell maturation and antigen-specific T-cell responses. Cancer Immunol Immunother 2020.

47. Schilderink R, Bell M, Reginato E et al. BET bromodomain inhibition reduces maturation and enhances tolerogenic properties of human and mouse dendritic cells. Mol Immunol 2016; 79: 66–76.

48. Toniolo PA, Liu S, Yeh JE et al. Inhibiting STAT5 by the BET bromodomain inhibitor JQ1 disrupts human dendritic cell maturation. J Immunol 2015; 194: 3180–3190.

49. Chee J, Wilson C, Buzzai A et al. Impaired T cell proliferation by ex vivo BET-inhibition impedes adoptive immunotherapy in a murine melanoma model. Epigenetics 2020; 15:134–144.

50. Gibbons HR, Mi DJ, Farley VM et al. Bromodomain inhibitor JQ1 reversibly blocks IFN-gamma production. Sci Rep 2019; 9: 10280.

51. Veneziani I, Fruci D, Compagnone M et al. The BET-bromodomain inhibitor JQ1 renders neuroblastoma cells more resistant to NK cell-mediated recognition and killing by downregulating ligands for NKG2D and DNAM-1 receptors. Oncotarget 2019; 10: 2151–2160.

52. Jespersen H, Olofsson Bagge R, Ullenhag G et al. Concomitant use of pembrolizumab and entinostat in adult patients with metastatic uveal melanoma (PEMDAC study): protocol for a multicenter phase II open label study. BMC Cancer 2019; 19: 415.

