## Supplementary figures and images for "Epigenetic therapy to enhance therapeutic effects of PD-1 inhibition in uveal melanoma"

### Supplementary Fig. S1

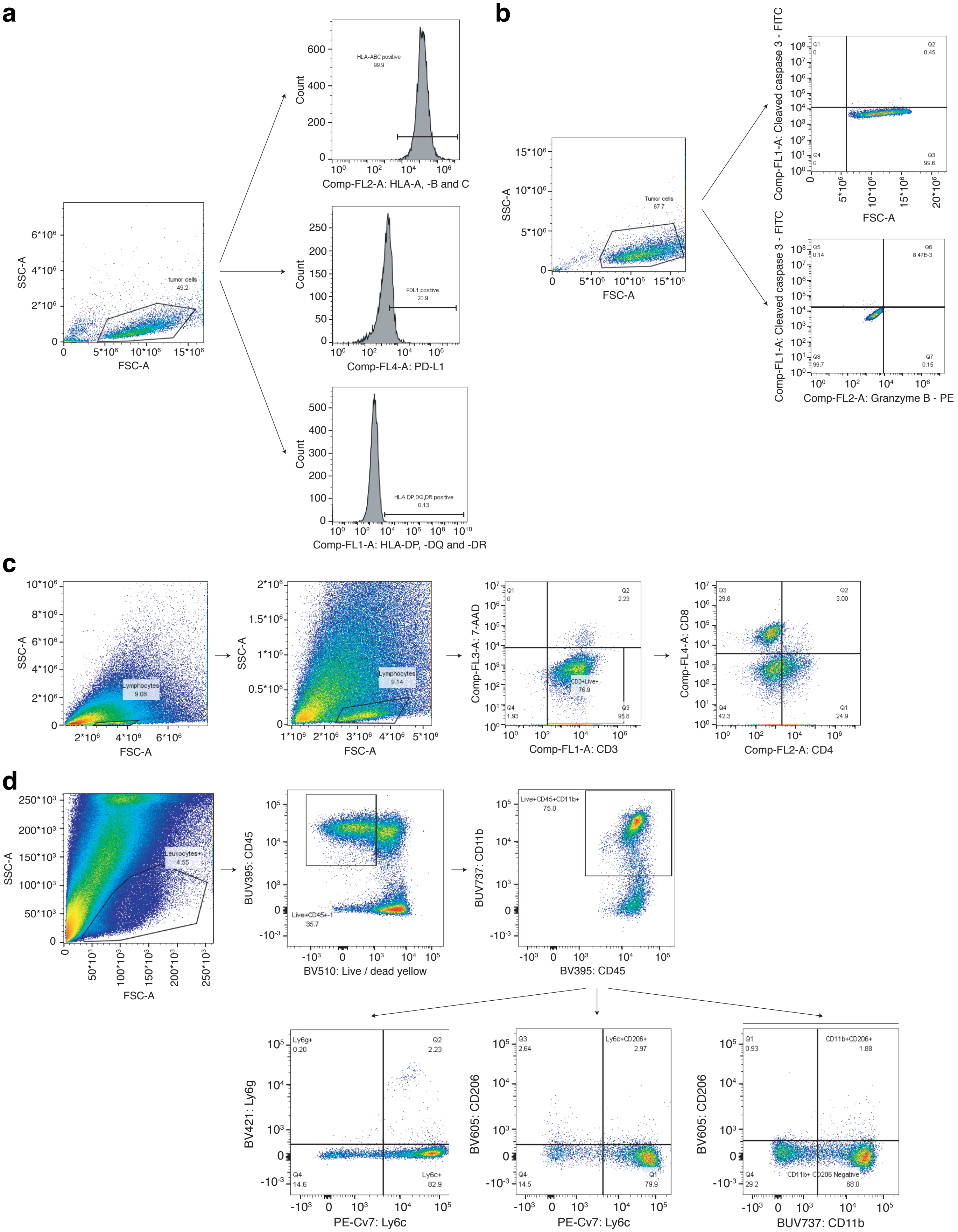

### Supplementary Fig. S2

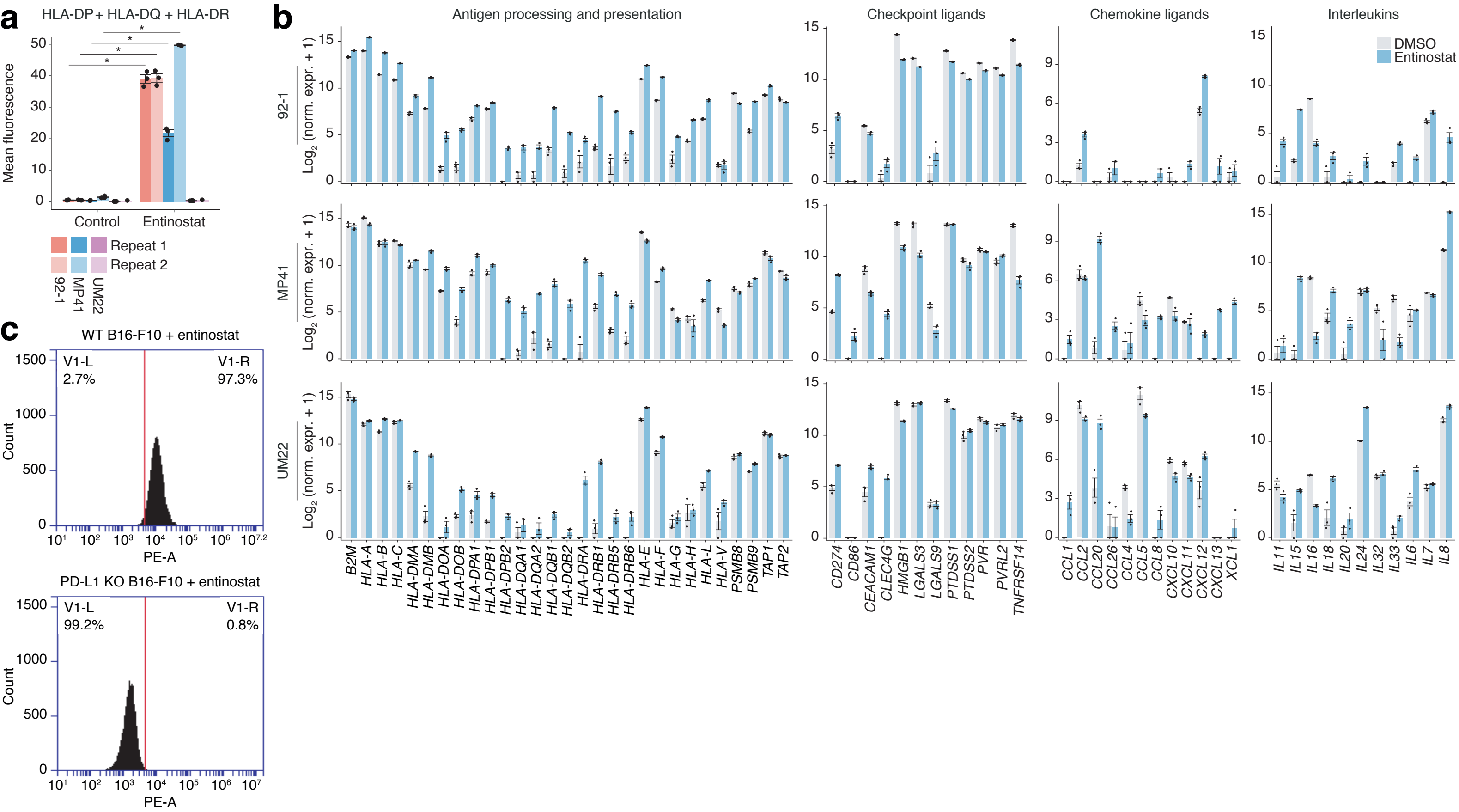
